# Microbiota epigenetically direct tuft cell differentiation to control type 2 immunity

**DOI:** 10.1101/2023.03.02.530792

**Authors:** Emily M. Eshleman, Taylor Rice, Crystal Potter, Amanda Waddell, Seika Hashimoto-Hill, Vivienne Woo, Sydney Field, Laura Engleman, Michael A. Schumacher, Mark R. Frey, Lee A. Denson, Fred D. Finkelman, Theresa Alenghat

## Abstract

Allergy and anti-helminth immunity are driven by type 2 responses in mucosal tissues. Tuft cells are key regulators of type 2 immunity, however the factors that control these cells remain poorly understood. Here we find that butyrate-producing commensal bacteria decrease tuft cells in the intestine. Butyrate suppression of tuft cells required the epigenetic modifying enzyme histone deacetylase 3 (HDAC3), suggesting that HDAC3 may promote tuft cell-dependent immunity. Consistent with this, epithelial-intrinsic HDAC3 actively regulated tuft cell expansion *in vivo* and was required to induce type 2 immune responses during helminth infection. Interestingly, butyrate epigenetically restricted stem cell differentiation into tuft cells, and inhibition of HDAC3 in adult mice and human intestinal organoids was sufficient to block tuft cell expansion. Collectively, these data reveal an epigenetic pathway in stem cells that directs tuft cell differentiation, and highlight a new level of regulation through which commensal bacteria calibrate intestinal immunity.

## INTRODUCTION

More than three billion individuals worldwide suffer from helminth infections or allergic diseases.^1^ Type 2 immunity is critical for host protection against pathogenic helminths. However, allergic disorders are induced and sustained by pathologic type 2 immune responses.^2^ Therefore, deciphering the mechanisms that direct type 2 immunity is needed to guide effective approaches to prevent or treat these highly prevalent conditions.

Tuft cells are secretory epithelial cells that are essential for initiating and maintaining type 2 immunity in the intestine. Furthermore, mice deficient in tuft cells are highly susceptible to helminth infection^3–5^ and models of inflammatory bowel disease.^6–8^ Intestinal tuft cells sense luminal signals and are exclusive producers of the type 2 cytokine, IL-25.^4^ Tuft cell derived-IL-25 activates immune cells, including CD4^+^ Th2 cells and type 2 innate lymphoid cells (ILC2s), to produce classical type 2 effector cytokines, ultimately leading to helminth clearance.^9–13^ In addition, the type 2 cytokine IL-13 stimulates changes to the intestinal epithelium, including tuft cell differentiation.^3–5^ Therefore, tuft cells drive a feedforward loop with local immune cells to promote their own lineage differentiation and anti-parasitic immunity, emphasizing the importance of tuft cells in sustaining type 2 immunity. While recent studies have begun to unravel the factors necessary for driving tuft cell hyperplastic response, the central regulatory mechanisms controlling tuft cell development and function remain poorly understood.

The trillions of microbes that reside in the intestinal tract, termed the microbiota, are essential for calibrating mammalian immune responses. Increasing evidence suggests that the intestinal microbiota exert strong regulatory effects on type 2 immune responses.^2, 14, 15^ Further, rising rates of inflammatory diseases and allergic reactions are associated with changes in microbiota composition and diversity.^14, 15^ Directly demonstrating a microbial influence, germ-free (GF) and antibiotic-treated mouse models exhibit increased type 2 immune responses characterized by elevated type 2 cytokines, increased type 2 effector cells, and enhanced susceptibility to anaphylaxis and allergic responses.^14, 15^ In addition, colonization of mice with microbiota from healthy children protected GF mice from allergic anaphylaxis, whereas this was not the case when mice were colonized with microbiota from food allergy patients^16^, suggesting that specific commensal microbes dampen type 2 immunity. While tuft cells are essential for controlling type 2 immunity, how the microbiota or microbiota-derived signals regulate tuft cells is not well understood.

Here we report that short-chain fatty acid (SCFA)-producing commensal bacteria limit tuft cell-induced type 2 immune responses. The SCFA butyrate suppressed IL-13-induced tuft cell hyperplasia in both mice and humans. Environmental cues derived from the microbiota can influence transcriptional programs of mammalian cells through epigenetic mechanisms.^17, 18^ Interestingly, butyrate constrained tuft cell development through inhibition of the microbial-sensitive epigenetic modifying enzyme, histone deacetylase 3 (HDAC3). Epithelial-intrinsic HDAC3 expression was required for activation of the tuft cell-ILC2 pathway and promoted pathogen clearance of the IL-13-sensitive helminth, *Nippostrongylus brasiliensis*. Furthermore, butyrate epigenetically restricted tuft cell differentiation, and targeted loss of HDAC3 within intestinal stem cells blocked tuft cell differentiation and downstream ILC2 responses. Taken together, these findings reveal an epigenetic pathway by which microbiota regulate tuft cell development and type 2 immune responses.

## RESULTS

### Tuft cell expansion in the intestine is suppressed by microbiota-derived butyrate

Succinate increases tuft cell numbers, promotes IL-25 expression, and activates type 2 innate lymphoid cells (ILC2s), in a tuft cell-dependent manner.^19–22^ Therefore, to determine how commensal microbes influence tuft cell-dependent type 2 immunity, succinate was administered to germ-free (GF) and conventionally-housed (CNV) microbiota-replete mice. Interestingly, CNV mice exhibited reduced tuft cell hyperplasia (**Figures 1A and 1B**) and decreased intestinal ILC2s relative to GF mice (**Figure 1C**), suggesting that the microbiota suppress tuft cell-dependent immune responses. To identify types of bacteria that may impact tuft cell responses, mice were treated with the antibiotic vancomycin to deplete Gram-positive bacterial species. Similar to GF mice, vancomycin-treatment increased intestinal tuft cells (**Figures 1D and 1E**) and ILC2s (**Figure 1F**) in response to succinate stimulation, suggesting that vancomycin-sensitive bacterial strains can inhibit tuft cell-dependent immunity. Vancomycin depletes *Clostridiales* bacterial families and reduces luminal concentrations of the SCFA, butyrate.^23^ Thus, to investigate whether SCFA-producing bacteria influence tuft cell-dependent immunity, GF mice were mono-associated with the butyrate producer *Faecalibacterium prausnitizii*. Remarkably, colonization with *F. prausnitizii* suppressed succinate-induced tuft cell hyperplasia (**Figures 1G and 1H**) and downstream ILC2s (**Figure 1I**) relative to GF mice. Gas-chromatography mass spectrometry analysis confirmed that mono-association with *F. prausnitizii* increased luminal concentration of butyrate compared to GF controls (**Figure 1J**). Taken together, these data indicate that butyrate-producing commensal bacteria can dampen tuft cell-dependent type 2 immunity in the intestine.

**Figure 1.**
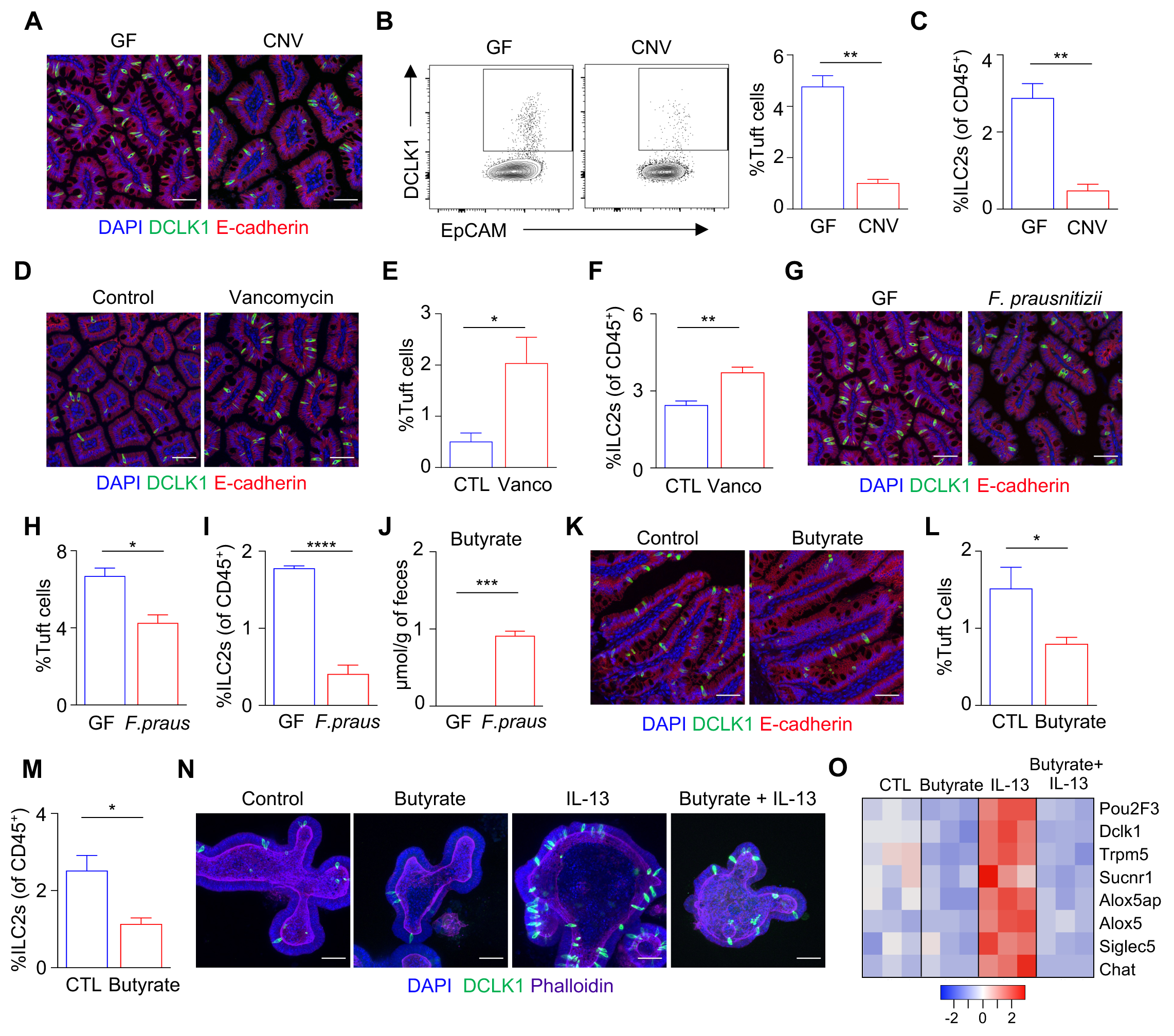
Tuft cell expansion in the intestine is suppressed by microbiota-derived butyrate. **(A)** Fluorescence staining of tuft cells (DCLK1^+^, green) in ileum, **(B)** frequency of DCLK1^+^ tuft cells and **(C)** ILC2s by flow cytometry in GF and CNV mice treated with succinate. **(D)** Fluorescence staining of tuft cells in ileum, **(E)** frequency of DCLK1^+^ tuft cells and **(F)** ILC2s by flow cytometry from control and vancomycin-treated mice receiving succinate. **(G)** Fluorescence staining of tuft cells in ileum, **(H)** frequency of DCLK1^+^ tuft cells and **(I)** ILC2s by flow cytometry from GF and *F. prausnitizii* mono-colonized mice treated with succinate. **(J)** Fecal concentration of butyrate in GF and *F. prausnitizii* mono-colonized mice. **(K)** Fluorescence staining of tuft cells in ileum, **(L)** frequency of DCLK1^+^ tuft cells, and **(M)** ILC2s by flow cytometry from WT mice treated with butyrate. **(N)** Fluorescence staining of tuft cells in organoids and **(O)** tuft cell-enriched gene expression by RNA-sequencing of WT organoids treated with butyrate, IL-13, or butyrate + IL-13. Scale bars, 50µM. ILC2s gated Live, CD45^+^, Lineage (CD4, CD8a, CD11b, CD11c, B220, Ly6G)^-^, CD90.2^+^, CD127^+^, Sca-1^+^, KLRG1^+^. Data are representative of at least three independent experiments for organoids and two independent mouse experiments, 3-5 mice per group. *p<0.05, **p<0.01, ***p<0.001, ****p<0.0001.

To directly test whether butyrate impacts tuft cell-dependent immunity, mice were supplemented with butyrate in their drinking water. Butyrate significantly reduced tuft cells (**Figures 1K and 1L**) and ILC2s (**Figure 1M**). To determine the epithelial-intrinsic effects of butyrate on tuft cell immune regulation, intestinal epithelial organoid cultures were derived from the intestine of wild-type mice. Intestinal organoids stimulated with the type 2 cytokine, IL-13, demonstrated increased tuft cells (**Figure 1N**). Interestingly, organoids treated with butyrate exhibited decreased basal tuft cell numbers and diminished IL-13- induced tuft cell hyperplasia (**Figure 1N**). In addition to protein characterization, global transcriptional analyses revealed that IL-13 induced tuft cell-associated genes in intestinal organoids, however butyrate blocked IL-13 induction of the tuft cell transcriptional signature (**Figure 1O**). Together, these data demonstrate that commensal microbial colonization, and specifically the microbiota-derived metabolite butyrate, can significantly shape tuft cell-dependent type 2 immunity.

### Butyrate controls tuft cell dynamics through histone deacetylase 3

Butyrate functions as a histone deacetylase (HDAC) inhibitor.^24, 25^ Furthermore, HDAC3 is a potent histone deacetylase^24, 26^ that incorporates microbiota-derived signals to maintain intestinal homeostasis.^25, 27^ Thus, to determine whether the butyrate-producer *F. prausnitizii* repressed succinate-induced tuft cell immunity by inhibition of epithelial HDAC3, HDAC3-sufficient (HDAC3^FF^) and IEC-deficient (HDAC3^ΔIEC^) mice derived under GF conditions were mono-associated with *F. prausnitizii*. *F. prausnitizii* reduced succinate-induced tuft cell hyperplasia (**Figures 2A and 2B**) and suppressed ILC2 responses (**Figure 2C**) in HDAC3^FF^, compared to GF HDAC3^FF^ controls. However, succinate was unable to promote tuft cell hyperplasia (**Figures 2A and 2B**) or expansion of intestinal ILC2s in GF-HDAC3^ΔIEC^ mice (**Figure 2C**). Furthermore, tuft cells and ILC2s were not affected in *F. prausnitizii* mono-colonized HDAC3^ΔIEC^ mice (**Figures 2A-2C**), supporting that *F. prausnitizii*-induced suppression of tuft cells is mediated by HDAC3. To next directly test whether butyrate regulates tuft cell development through HDAC3, organoids were derived from HDAC3^FF^ and inducible villin-cre mice (HDAC3^ΔIEC-IND^) (**Figure 2D**), in which tamoxifen treatment significantly decreased *Hdac3* expression in HDAC3^ΔIEC-IND^ organoids, but not tamoxifen treated HDAC3^FF^ controls (**Figure 2E**). Butyrate inhibited tuft cell-associated genes including the critical transcription factor, *Pou2f3*, and tuft cell marker, *Dclk1*, in control organoids (**Figures 2F and 2G**). Consistent with this expression profile, tuft cell numbers were reduced in butyrate treated control organoids (**Figure 2H**). Loss of HDAC3 reduced basal tuft cell gene expression and tuft cell numbers (**Figures 2F-2H**). In addition, butyrate failed to further suppress tuft cells in HDAC3-depleted organoids (**Figures 2F-2H**), supporting that butyrate suppresses tuft cells via HDAC3 inhibition, and that HDAC3 is required for tuft cell homeostasis.

**Figure 2.**
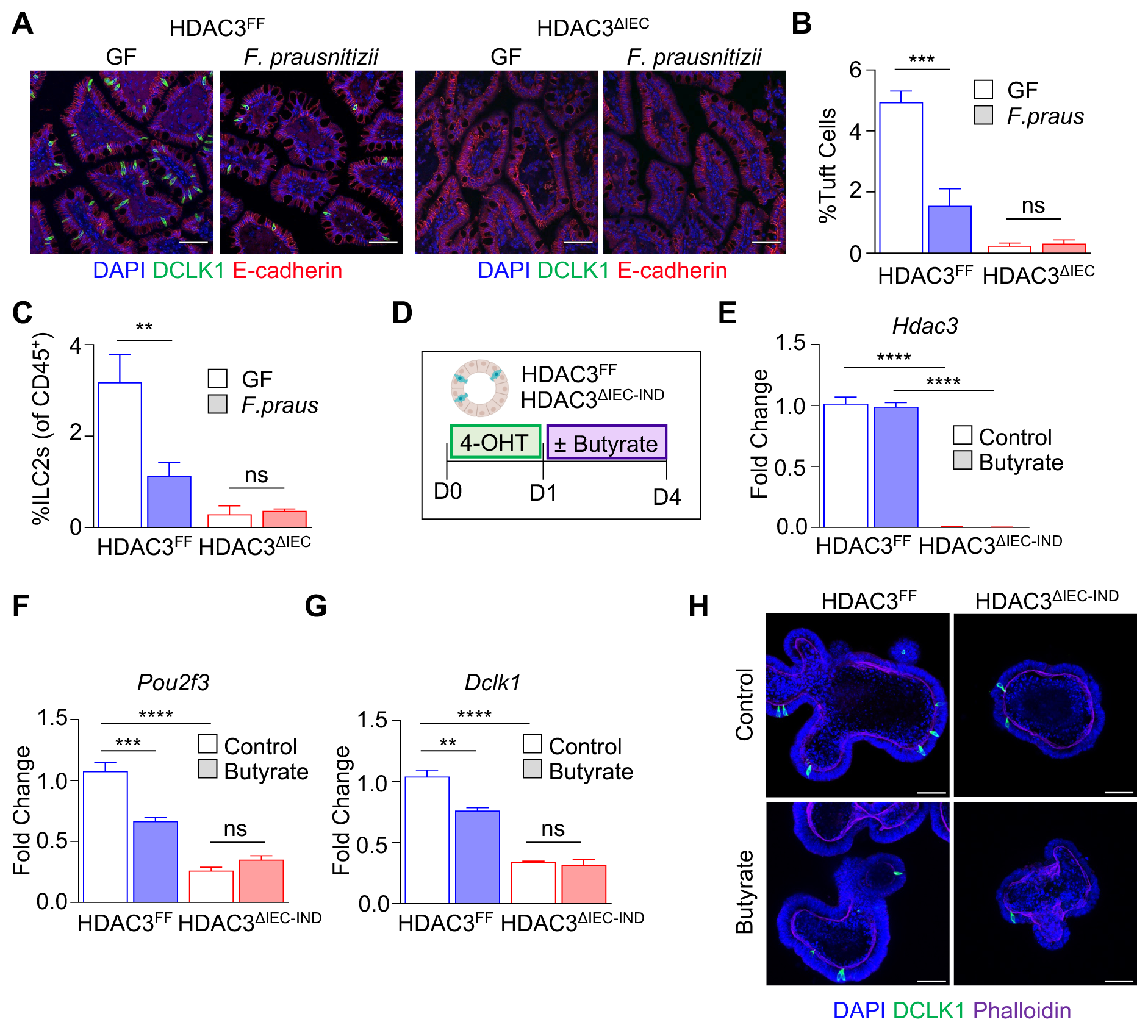
Butyrate controls tuft cell dynamics through histone deacetylase 3. **(A)** Fluorescence staining of tuft cells (DCLK1^+^, green) in ileum, **(B)** frequency of DCLK1^+^ tuft cells and **(C)** ILC2s by flow cytometry in GF and *F. prausnitizii* mono-associated HDAC3^FF^ and HDAC3^ΔIEC^ mice stimulated with succinate. ILC2s gated Live, CD45^+^, Lineage (CD4, CD8a, CD11b, CD11c, B220, Ly6G)^-^, CD90.2^+^, CD127^+^, Sca-1^+^, KLRG1^+^. **(D)** Experimental diagram. **(E)** mRNA expression of *Hdac3*, **(F)** *Pou2f3*, and **(G)** *Dclk1*, and **(H)** fluorescence staining of tuft cells in organoids treated with tamoxifen (4-OHT) +/- butyrate. Scale bars, 50µM. Data are representative of at least three independent experiments for organoids and two independent mouse experiments, 3-4 mice per group. *p<0.05, **p<0.01, ***p<0.001, ****p<0.0001, ns=not significant.

### Epithelial HDAC3 directs intestinal type 2 immunity and effective worm clearance

Tuft cells are essential for initiating and sustaining type 2 immune responses to helminth infection and promoting worm clearance.^3–5^ Consistent with this work, we found that infection with *Nippostrongylus brasiliensis* enhanced tuft cell hyperplasia (**Figures 3A and 3B**), epithelial *IL-25* expression, and intestinal ILC2s (**Figures 3C and 3D**). Remarkably, loss of IEC-intrinsic HDAC3 expression resulted in elevated intestinal *N. brasiliensis* egg counts (**Figure 3E**). Furthermore, several HDAC3^FF^ mice cleared the infection by day 10, while HDAC3^ΔIEC^ mice retained high worm burdens (**Figure 3F**), indicating that epithelial HDAC3 promotes effective clearance. Intestinal tuft cell numbers increased in response to *N. brasiliensis* infection in control mice (**Figures 3G and 3H**). However, *N. brasiliensis-*infected HDAC3^ΔIEC^ mice failed to demonstrate this tuft cell expansion (**Figures 3G and 3H**). Consistent with a lack of helminth-induced tuft cell hyperplasia, downstream *IL-25* (**Figure 3I**), ILC2s (**Figure 3J**), and IL-13 responses were impaired in HDAC3^ΔIEC^ mice (**Figure 3K**). Collectively, these data support that epithelial expression of HDAC3 is necessary for effective type 2-driven tuft cell development.

**Figure 3.**
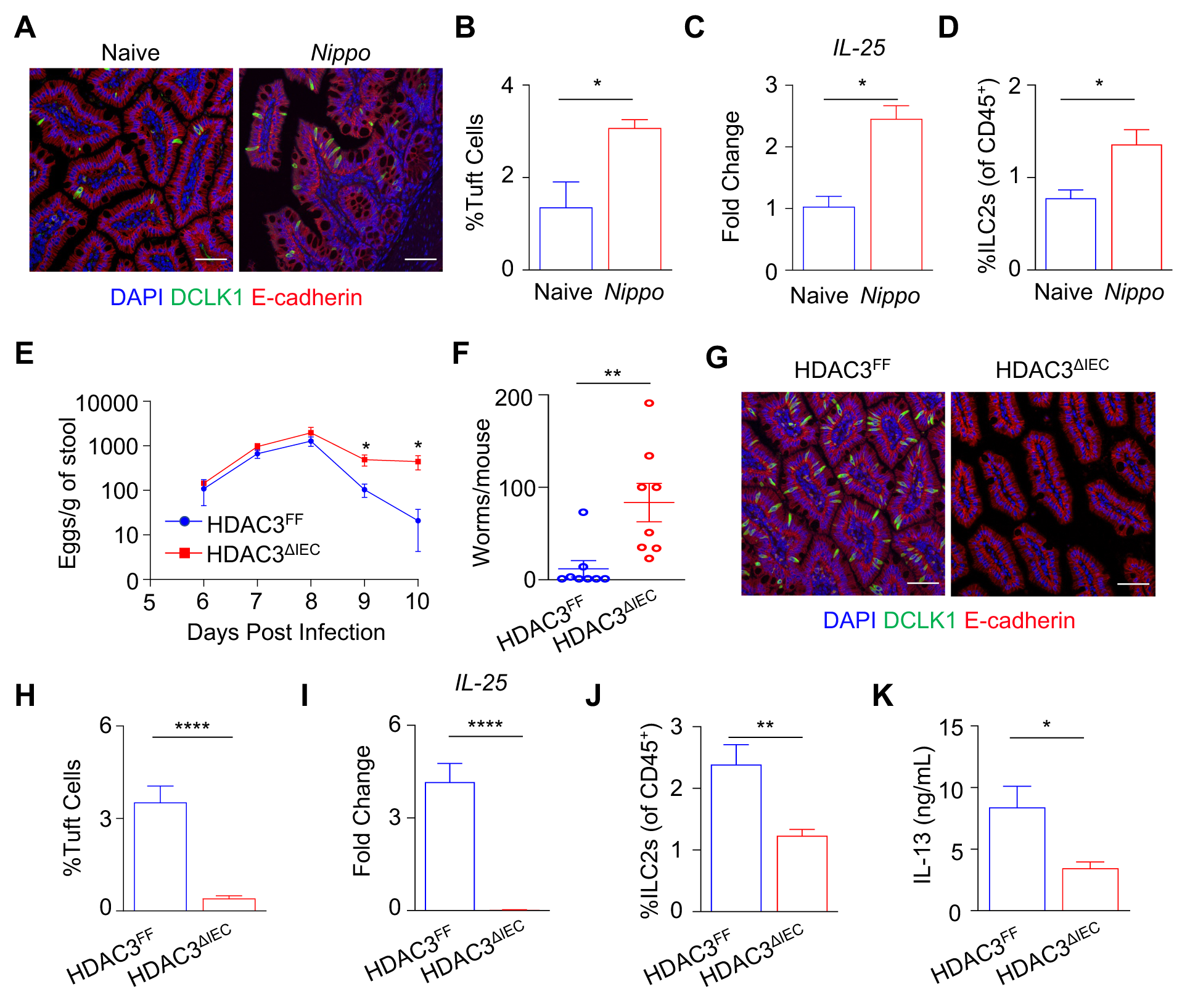
Epithelial HDAC3 directs intestinal type 2 immunity and effective worm clearance. **(A)** Fluorescence staining of tuft cells (DCLK1^+^, green), **(B)** frequency of DCLK1^+^ tuft cells by flow cytometry, **(C)** mRNA expression of *IL-25* in SI IECs, and **(D)** frequency of ILC2s in ileum of WT mice naïve and 7 days post-*N. brasiliensis* infection. **(E)** Fecal *N. brasiliensis* egg counts and **(F)** intestinal *N. brasiliensis* worm counts at day 10 in HDAC3^FF^ and HDAC3^ΔIEC^ mice infected with *N. brasiliensis*. **(G)** Fluorescence staining of tuft cells in ileum, **(H)** frequency of DCLK1^+^ tuft cells by flow cytometry, **(I)** mRNA expression of *IL-25* from SI IECs, **(J)** frequency of lamina propria ILC2s, and **(K)** serum concentration of IL-13 in HDAC3^FF^ and HDAC3^ΔIEC^ mice 10 days post *N. brasiliensis* infection. ILC2s gated Live, CD45^+^, lineage (CD4, CD8a, CD11b, CD11c, B220, Ly6G)^-^, CD90.2^+^, CD127^+^, Sca-1^+^, KLRG1^+^. Scale bars, 50µM. Data are representative from at least two independent experiments, 4-8 mice per group, per timepoint. *p<0.05, **p<0.01, ***p<0.001, ****p<0.0001.

### Active regulation by HDAC3 is necessary for IL-13-induced tuft cell expansion

To investigate whether expansion of tuft cells in the adult intestine reflect active regulation by HDAC3 *in vivo*, HDAC3^ΔIEC-IND^ mice were supplemented with succinate prior to HDAC3 loss to induce tuft cell hyperplasia, and then treated with either vehicle or tamoxifen. Mice that received vehicle-treatment displayed elevated succinate-induced tuft cell numbers (**Figure 4A and 4B**) and ILC2 responses (**Figure 4C**). However, mice treated with tamoxifen to delete epithelial HDAC3 exhibited complete loss of tuft cells and diminished ILC2s responses (**Figure 4A-4C**), indicating that active regulation by HDAC3 *in vivo* was required for tuft cell hyperplasia. To determine whether epithelial HDAC3 regulation of tuft cells affects responsiveness to IL-13, intestinal organoids were generated from HDAC3^FF^ and HDAC3^ΔIEC-IND^ mice and treated with tamoxifen prior to IL-13 stimulation. Importantly, IL-13 did not affect *Hdac3* expression (**Figure 4D**). HDAC3-sufficient organoids rapidly expanded tuft cells in response to IL-13 stimulation, however loss of HDAC3 blocked IL- 13-induced tuft cell hyperplasia (**Figures 4E and 4F**). Consistently, loss of epithelial HDAC3 inhibited induction of tuft cell genes, including *Pou2f3* and *Dclk1*, in response to IL-13 (**Figures 4G and 4H**). Therefore, these data reveal that epithelial HDAC3 actively directs the tuft cell response to IL-13.

**Figure 4.**
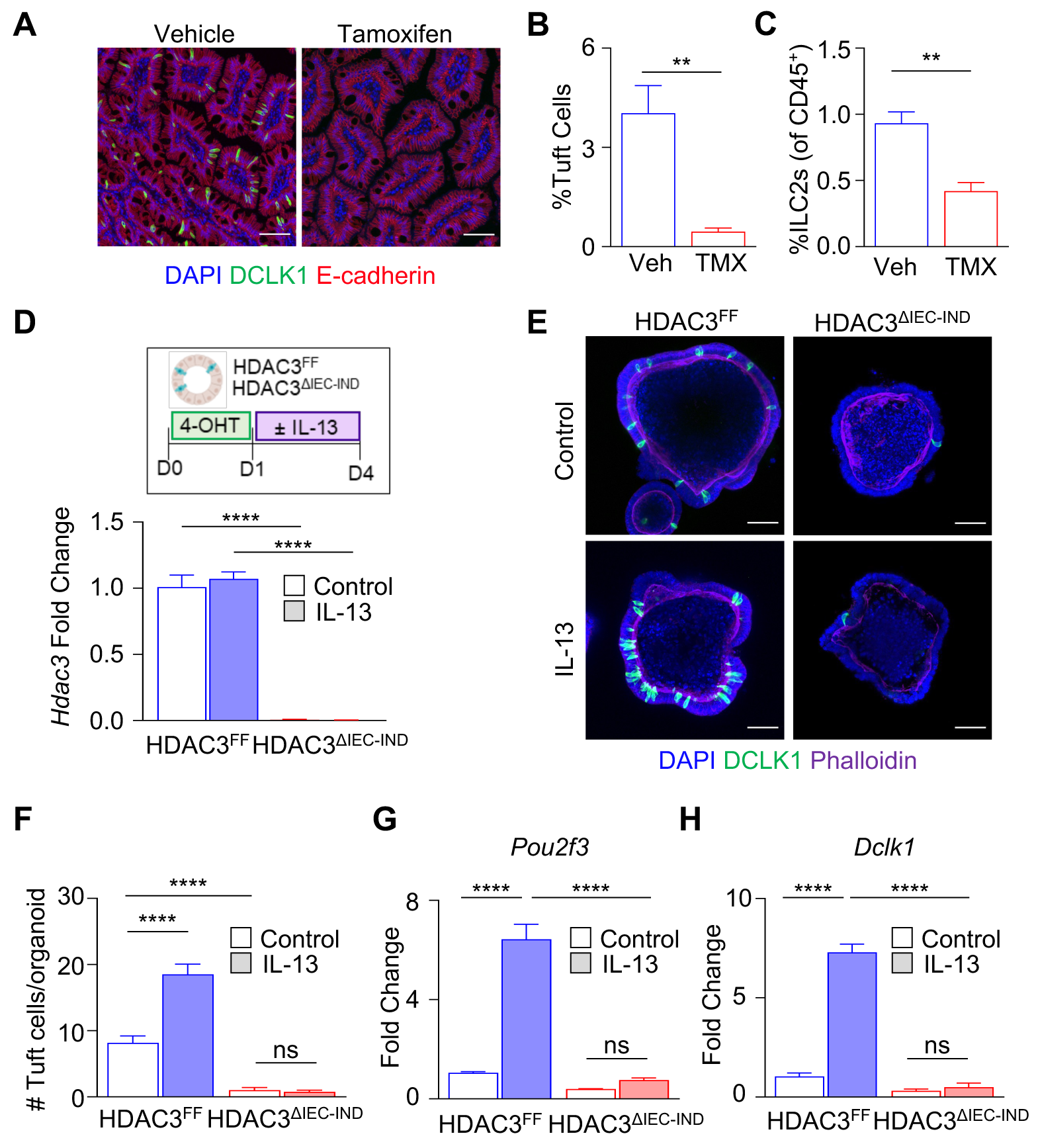
Active regulation by HDAC3 is necessary for IL-13-induced tuft cell expansion. **(A)** Fluorescence staining of tuft cells (DCLK1^+^, green) in ileum, **(B)** frequency of DCLK1^+^ tuft cells and (**C**) ILC2s by flow cytometry in HDAC3^ΔIEC-IND^ mice treated with succinate then vehicle or tamoxifen. ILC2s gated Live, CD45^+^, lineage (CD4, CD8a, CD11b, CD11c, B220, Ly6G)^-^, CD90.2^+^, CD127^+^, Sca-1^+^, KLRG1^+^. **(D)** mRNA expression of *Hdac3* in HDAC3^FF^ and HDAC3^ΔIEC-IND^ organoids treated with tamoxifen (4-OHT) +/- IL-13. (**E**) Fluorescence staining of tuft cells in organoids, (**F**) quantification of DCLK1^+^ tuft cells per organoid, (**G**) *Pou2f3* and (**H**) *Dclk1* mRNA expression in HDAC3^FF^ and HDAC3^ΔIEC-IND^ organoids treated with 4-OHT +/- IL-13. Scale bars, 50µM. Data are representative from at least three independent experiments, 3-4 mice per group. *p<0.05, **p<0.01, ***p<0.001, ****p<0.0001, ns=not significant.

### Human tuft cell differentiation is restricted by butyrate inhibition of HDAC3

Our understanding of human tuft cell differentiation remains largely unknown. Thus, to next directly examine regulation of human tuft cells, we generated organoids from ileal endoscopic biopsies harvested from healthy non-IBD patients. IL-13 exposure promotes tuft cell differentiation in murine intestinal organoids, however this has not previously been shown for human-derived organoids. Basal tuft cells in ileal-derived human organoids were generally undetectable by immunofluorescence staining for the human tuft cell marker, p-EGFR (**Figure 5A**). Interestingly, IL-13 promoted tuft cell development in human intestinal organoids derived from multiple different patients (**Figure 5A**), suggesting similarities between human and murine tuft cell responses. Furthermore, butyrate blunted IL-13-induced tuft cell genes including *POU2F3* and the tuft cell marker, *TRPM5* (**Figures 5B and 5C**) and suppressed tuft cell responses to IL-13 in human organoids (**Figure 5D**). In addition, treatment with an HDAC3-specific inhibitor (HDAC3i: RGFP966) similarly reduced IL-13-induced tuft cell genes (**Figures 5E and 5F**) and tuft cell development in human organoids (**Figure 5G**). Butyrate treatment in the presence of HDAC3 inhibition did not further block tuft cells (**Figures 5H and 5I**). Therefore, similar to our murine findings, butyrate controls tuft cell differentiation via HDAC3 in human intestine. Collectively, these data demonstrate that active regulation by HDAC3 controls IL-13-induced tuft cell hyperplastic responses in both mice and humans, and suggests a central role for HDAC3 in tuft cell progenitor cells.

**Figure 5.**
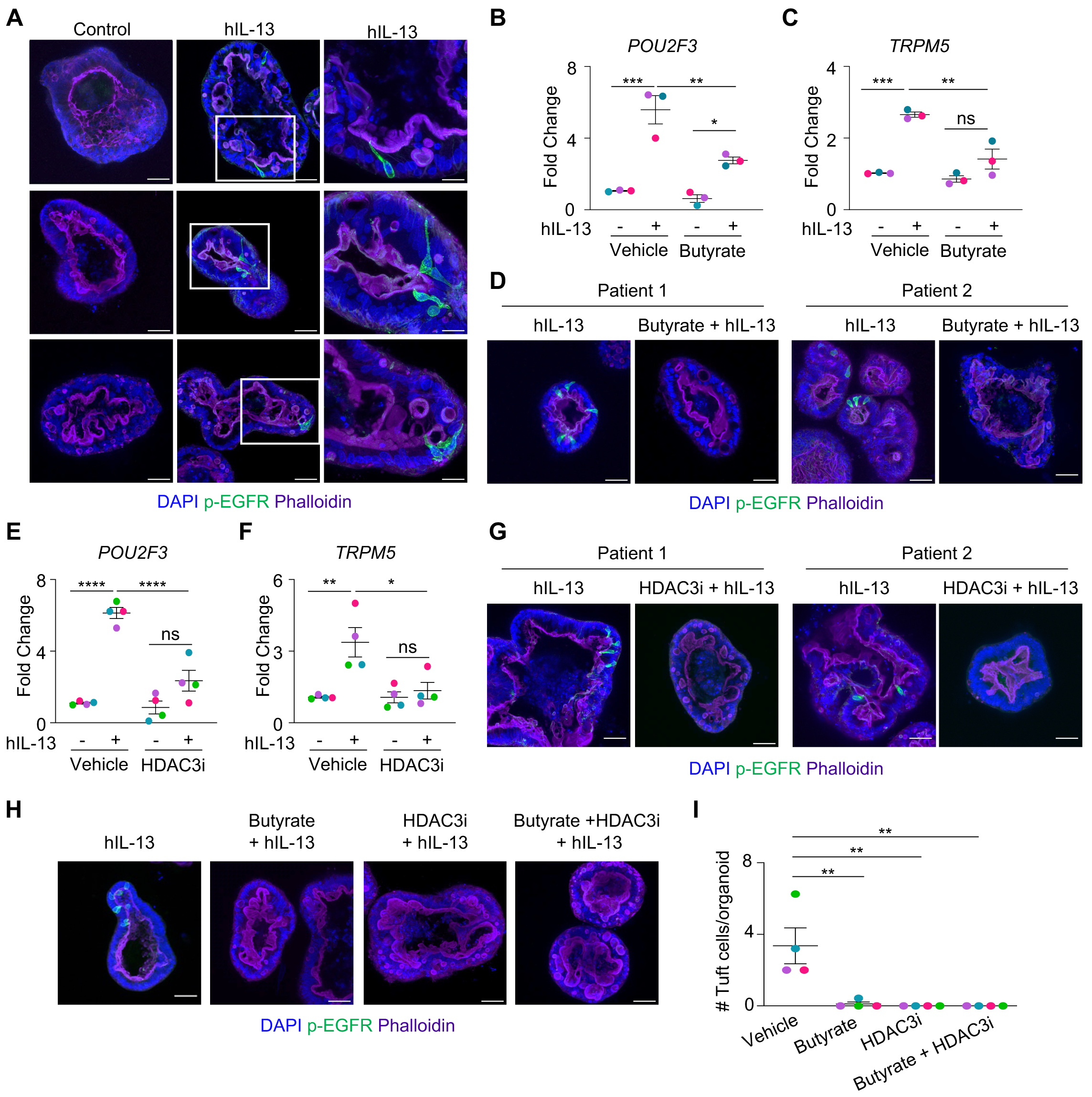
Human tuft cell differentiation is restricted by butyrate inhibition of HDAC3. **(A)** Fluorescence staining of tuft cells (p-EGFR^+^, green) in human crypt-derived intestinal organoids treated with IL-13. Scale bars, 50µM or 20µM for in-lay images. **(B)** mRNA expression of *POU2F3* and **(C)** mRNA expression of *TRPM5* in organoids treated with IL-13 +/- butyrate, n=3 patients. **(D)** Fluorescence staining of tuft cells in human intestinal organoids treated with IL-13 +/- butyrate. **(E)** mRNA expression of *POU2F3* and **(F)** mRNA expression of *TRPM5* in organoids treated with IL-13 +/- HDAC3i (RGFP966), n=4 patients. **(G)** Fluorescence staining of tuft cells in human intestinal organoids treated with IL-13 +/- HDAC3i (RGFP966). **(H)** Fluorescence staining of tuft cells in human intestinal organoids treated with IL-13 +/- butyrate and/or HDAC3i. **(I)** Quantification of human p-EGFR^+^ tuft cells, n=4 patients. Data are from organoids derived from 3-4 different patients, each color in the panel represents distinct patients. *p<0.05, **p<0.01, ***p<0.001, ****p<0.0001, ns=not significant.

### Stem cell-intrinsic HDAC3 epigenetically promotes tuft cell differentiation

Recent work identified that Sprouty2 (Spry2), a negative regulator of MAP kinase signal transduction, suppresses tuft cell development.^28^ Therefore, we generated mice lacking Spry2 specifically in intestinal epithelial cells by breeding floxed Spry2 mice (Spry2^FF^) to mice expressing Cre recombinase downstream of the villin promoter (Spry2^ΔIEC^). Consistent with the role of Spry2 in inhibition of tuft cell expansion in the colon^28^, these mice also demonstrated increased tuft cells in the ileum (**Figures 6A and 6B**). Interestingly, histone H3K9 acetylation (H3K9ac) was elevated at the *Spry2* promoter in HDAC3^ΔIEC^ cells (**Figure 6C**), suggesting Spry2 may be a direct target of HDAC3. Small intestinal crypt cells from HDAC3^ΔIEC^ mice exhibited increased *Spry2* expression relative to controls (**Figure 6D**), supporting that HDAC3 epigenetically represses this suppressor of tuft cell differentiation.

**Figure 6.**
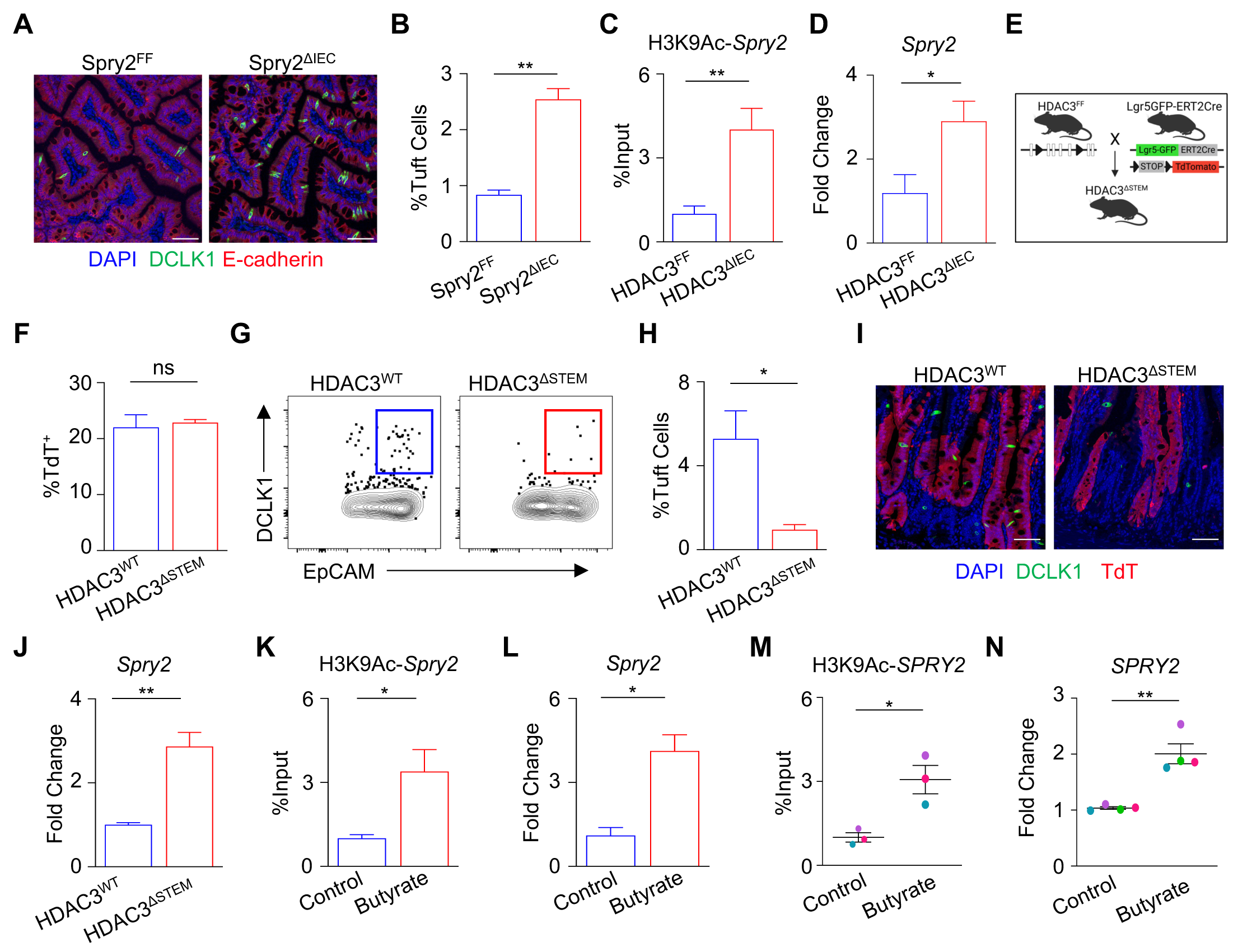
Stem cell-intrinsic HDAC3 epigenetically promotes tuft cell differentiation. **(A)** Fluorescence staining of tuft cells (DCLK1^+^, green) and **(B)** frequency of DCLK1^+^ tuft cells by flow cytometry in ileum of Spry2^FF^ and Spry2^ΔIEC^. **(C)** Chip-qPCR for H3K9ac in *Spry2* promoter in HDAC3^FF^ and HDAC3^ΔIEC^ IECs. **(D)** mRNA expression of *Spry2* in HDAC3^FF^ and HDAC3^ΔIEC^ SI crypt-associated cells. **(E)** Diagram of HDAC3^ΔSTEM^ mice. **(F)** Frequency of TdTomato^+^ (TdT) cells in EpCAM^+^ gate and **(G, H)** Frequency of DCLK1^+^ tuft cells from TdT^+^ gated cells in ileum of HDAC3^WT^ and HDAC3^ΔSTEM^ mice. **(I)** Fluorescence staining of tuft cells in ileum in HDAC3^WT^ and HDAC3^ΔSTEM^ mice. **(J)** mRNA expression of *Spry2* in HDAC3^WT^ and HDAC3^ΔSTEM^ SI crypt-associated cells. **(K)** Chip- qPCR for H3K9ac in *Spry2* promoter and **(L)** mRNA expression of *Spry2* in intestinal organoids stimulated with butyrate. **(M)** Chip-qPCR for H3K9ac at hSprouty2 (*SPRY2*) promoter in human crypt-derived intestinal organoids treated +/- butyrate, n=3 patients, each color in the panel represents distinct patients. **(N)** mRNA expression of *SPRY2* in human intestinal organoids +/- butyrate, n=4 patients, each color in the panel represents distinct patients. Scale bars, 50µM. Data are representative from at least three independent experiments, 3-4 mice per group. *p<0.05, **p<0.01, ***p<0.001, ****p<0.0001, ns=not significant.

Lgr5^+^ intestinal epithelial stem cells (ISCs) continuously give rise to all differentiated IEC lineages, including tuft cells.^4^ To determine whether ISC expression of HDAC3 is required for tuft cell differentiation, mice were generated to specifically delete HDAC3 in Lgr5^+^- expressing ISCs (HDAC3^ΔSTEM^) with lineage-tracing TdTomato (TdT) downstream of a stop-flox codon (**Figure 6E**). Deletion of HDAC3 in Lgr5^+^ ISCs did not impact the overall frequency of TdT^+^ expressing cells (**Figure 6F**), supporting that IEC differentiation is not globally affected by loss of ISC-intrinsic HDAC3. However, loss of HDAC3 in Lgr5^+^ ISCs blunted baseline tuft cell numbers (**Figures 6G-6I**), indicating that ISC-intrinsic HDAC3 promotes tuft cell lineage differentiation from ISCs. Further, when supplemented with succinate, tamoxifen-treated HDAC3^ΔSTEM^ mice had impaired tuft cell differentiation, compared to vehicle treated controls (**Figure S1A and S1B**) and reduced ILC2s (**Figure S1C**), highlighting that stem cell-intrinsic HDAC3 regulation is necessary for inducing tuft cell differentiation. Interestingly, *Spry2* gene expression was elevated in crypt cells derived from HDAC3-deficient stem cells, relative to HDAC3-sufficient cells (**Figure 6J**). In addition, intestinal organoids stimulated with butyrate displayed H3K9ac enrichment at the *Spry2* promoter (**Figure 6K**) and increased *Spry2* gene expression (**Figure 6L**). Taken together, these data suggest that butyrate downregulates tuft cell differentiation by limiting HDAC3 activity at the *Spry2* promoter. Remarkably, butyrate also led to H3K9Ac enrichment at the *SPRY2* promoter in human intestinal organoids (**Figure 6M**), along with increased *SPRY2* expression (**Figure 6N**). Thus, functional pathways are established in the human intestine by which microbiota-derived metabolites can epigenetically direct human tuft cell differentiation.

## DISCUSSION

In this study, we identified a new pathway by which the microbiota calibrate intestinal tuft cell differentiation (**Figure S2**). Microbiota-derived butyrate limited IL-13-induced tuft cell hyperplasia and effective type 2 immune responses. HDAC3 specifically in stem cells epigenetically altered the tuft cell suppressor gene, *Spry2*, and promoted tuft cell differentiation. Remarkably, this level of microbiota-host regulation also controlled tuft cell development in the human intestine, highlighting essential clinical implications for either limiting or promoting type 2 immunity. Tuft cell differentiation requires expression of the taste-cell specific transcription factor, Pou2F3.^3, 29–31^ Recently, Pou2F3 co-activators have been identified and specific isoform usage has been suggested to modulate homeostatic tuft cell development.^32, 33^ The secretory lineage factors, ATOH1 and Sox4, are only partially required for tuft cell development, as deletion of either molecule results in partial or region-specific reduction in tuft cells.^34–37^ Here we find that stem cell-intrinsic loss of HDAC3 inhibited tuft cell differentiation. Interestingly, neither succinate nor IL-13 could overcome HDAC3 deficiency to promote tuft cell differentiation or expansion. Therefore, these data highlight that HDAC3 regulation of stem cells is essential for the tuft cell lineage program (**Figure S2**).

Canonically, HDAC3 deacetylates histone targets to alter gene expression and cellular functions. HDAC3 classically suppresses gene expression of direct targets via histone deacetylation. Loss of ISC-intrinsic HDAC3 resulted in increased histone acetylation and elevated gene expression of the tuft cell suppressor, *Spry2*. Previous work demonstrated that basal Spry2 directly inhibits PI3K/Akt signaling, which allows GSK3β control of tuft cell differentiation.^28, 38, 39^ Our data indicate that HDAC3 inhibition increases *Spry2* expression and histone acetylation, suggesting that HDAC3 may directly repress *Spry2* transcription. Consistent with HDAC3 inhibition, butyrate also enhanced *Spry2* expression and histone acetylation, supporting that butyrate mechanistically limits tuft cell differentiation in part by decreasing HDAC3 activity at the *Spry2* gene. While these data highlight one target through which butyrate can epigenetically restrict tuft cell differentiation, HDAC3 regulates several pathways to collectively alter gene expression and cellular activity. Thus, it is likely that multiple HDAC3-dependent mechanisms mediate butyrate-dependent regulation of tuft cell differentiation.

Butyrate has been shown to exert broad immunoregulatory effects and influence the development and function of the intestinal epithelium via SCFA-sensing receptors such as FFAR2, FFAR3, and GPR109A, or direct inhibition of HDAC activity.^40^ Previous studies have suggested that butyrate or other SCFAs, including propionate and acetate, may have little impact on tuft cell homeostasis in mouse models.^19, 21^ However, the intestinal crypt structure has been suggested to limit stem cell exposure to butyrate and other microbiota-derived factors.^41^ In organoids where crypt structure is diminished, butyrate inhibited tuft cell differentiation in both murine and human models. Beyond production of butyrate, the microbiota generate numerous other metabolites and factors that may positively regulate ISC function and tuft cell differentiation. Consistent with this, microbial-derived succinate promotes tuft cell differentiation. Thus, diverse microbial-derived signals are likely to synergistically or antagonistically fine-tune tuft cell development.

Microbiota composition has been linked with the development of several chronic inflammatory diseases including allergy, asthma, and IBD. The findings described here reveal a central role for commensal bacterial-derived metabolites in epigenetically limiting type 2 intestinal immune responses through active regulation of tuft cell differentiation. In addition to the intestine, tuft cells have been identified in the lung, trachea, esophagus, stomach, thymus, pancreas, and biliary tract, and contribute to airway hypersensitivity.^42–48^ Interestingly, HDAC3 is expressed ubiquitously, and butyrate levels at extra-intestinal sites reflect microbiota colonization. Thus, it is plausible that butyrate regulation of HDAC3 may control tuft cell-dependent immune responses at distant tissue sites as well. Therefore, modulating this pathway, pharmacologically or through diet or microbiota-based approaches, might be useful for treating pathologic inflammation across mucosal tissues.

## Acknowledgements

We thank members of the Center for Inflammation and Tolerance at CCHMC for useful discussions and critical reading of the manuscript. We thank CCHMC Veterinary Services, Research Flow Cytometry Core, Confocal Imaging Core, Pluripotent Stem Cell Facility, Pathology Research Core, Gene Expression Core, University of Cincinnati Genomics Core, and University of Louisville CREAM Core for services and technical assistance. This research is supported by the National Institutes of Health (DK114123, DK116868 to T.A.; DK095004, DK119694 to M.R.F; K01DK131390 to M.A.S.; and F32AI147591 to E.M.E.), and a Kenneth Rainin Foundation award to T.A. T.A. holds an Investigator in the Pathogenesis of Infectious Disease Award from the Burroughs Wellcome Fund. This project is supported in part by PHS grant P30 DK078392 and the Center for Stem Cell & Organoid Medicine at CCHMC.

## Author Contributions

E.M.E., T.R., S.F., C.P., V.W., S.H.H., A.W., L.E., F.D.F., and T.A. designed and performed the studies. E.M.E., T.R., and C.P. analyzed the data. M.A.S and M.R.F. provided expertise and mice. L.A.D. provided patient samples and clinical expertise. T.A. and E.M.E. wrote the manuscript. All authors edited the manuscript.

## METHODS

### Mice

Conventionally-housed C57BL/6J mice were purchased from Jackson Laboratories and maintained in our specific-pathogen free colony at Cincinnati Children’s Hospital Medical Center (CCHMC). Germ-free (GF) mice were maintained in flexible isolators in the CCHMC Gnotobiotic Mouse Facility, fed autoclaved feed and water, and monitored for absence of microbes. HDAC3^FF^ were generated as previously described^49^, and crossed to C57BL/6J-Villin-Cre (HDAC3^ΔIEC^)^50^, tamoxifen-inducible Villin-Cre mice (HDAC3^ΔIEC-IND^)^51^, and tamoxifen-inducible Lgr5GFP-ERT2Cre (HDAC3^ΔSTEM^)^52^ mice. Spry2 floxed mice (MMRRC 11469) were crossed to C57BL/6J-Villin-Cre to generate Spry2^ΔIEC^ mice as previously described.^28^ Sex-and age-matched littermate controls were used for all studies. Mice were housed up to 4-per cage in a ventilated cage system in a 12-h light/dark cycle, with free access to water and food. All mouse studies were conducted with approval by Animal Care and Use Committees at CCHMC. These protocols follow standards enacted by the United States Public Health Services and Department of Agriculture. All experiments followed standards set forth by Animal Research: Reporting of In Vivo Experiments (ARRIVE).

### GF mono-association and murine type 2 immune models

For tamoxifen-inducible models, mice were given daily intraperitoneal injections of 1 mg of tamoxifen (Sigma) for 5 days. Mice were provided 150 mM of sodium succinate hexahydrate (Alfa Aesar), 150 mM sodium butyrate (Sigma), or 0.5 mg/mL of vancomycin (GoldBio) dissolved in drinking water for 7-15 days. *Faecalibacterium prausnitizii* was cultured in YBHI Media (Brain Heart infusion media, 0.5% yeast extract, 5 mg/L Hemin, 1 mg/mL cellulose, 1 mg/mL maltose, 0.5 mg/mL) under anaerobic conditions as previously described.^25, 53^ For mono-association, GF mice were pre-treated with 0.2 M sodium bicarbonate for 10 mins prior to being gavaged with 2 x 10^9^ *F. prausnitizii* as previously described.^25, 53^ Mice were colonized for at least 14 days and mono-association was confirmed by 16s qPCR analysis. For 16s analysis, fecal DNA was isolated using QIAamp Fast DNA Stool Mini Kit (Qiagen) following the kit protocol and bacterial DNA was assessed by quantitative PCR (QuantStudio3; Applied Biosystems) using bacterial-specific or 16s primer pairs. For *Nippostrongylus brasiliensis* infection, mice were injected subcutaneously with 500 stage 3 larvae as previously described^54, 55^ and monitored daily for egg counts. Mice were sacrificed at day 7 and day 10 post-infection.

### Cell Isolation

The small intestine was harvested, opened, and washed in PBS. For IECs, tissue was placed in pre-warmed strip buffer (PBS, 5% FBS, 1 mM EDTA, 1 mM DTT) and incubated at 37°C at a 45-degree angle with shaking at 180-rpm for 15-min. For lamina propria isolation, tissue was washed with PBS to remove EDTA and DTT then incubated in pre-warmed digestion buffer (RPMI with 1-mg/mL collagenase/dispase (Sigma)) at 37°C at a 45-degree angle with shaking at 180-rpm for 30-min. After incubation, the tissue was vortexed and passed through a 70µM cell strainer.

### Flow cytometry

Cells were stained for flow cytometry using the following antibodies diluted in FACS buffer (2% FBS, 0.01% sodium azide, PBS): BV711-anti-CD326 (EpCAM) (Clone:G8.8, BD Bioscience), BUV395-anti-CD45.2 (Clone:104, BD Bioscience), APC-anti-GATA-3 (Clone: 16E10A23, Invitrogen), APC-eFluor-780-anti-KLRG-1 (Clone: 2F1, Invitrogen), Pe-Cy7-anti-CD127 (Clone: A7R34, Invitrogen), PE-CF594-anti-CD25 (Clone: PC61, BioLegend), PE-anti-SiglecF (Clone: E50-2440, BD Bioscience), PerCP-eFluor710-anti-CD4 (Clone: RM4-5, Invitrogen), PerCP-eFluor710-anti-CD8a (Clone: 53-6.7, Invitrogen), PerCP-eFluor710-anti-B220 (Clone: RA3-6B2, Invitrogen), PerCP-eFluor710-anti-Ly6G (Clone: 1A8-Ly6g, Invitrogen), PerCP-Cy5.5-anti-CD11b (Clone: M1/70, Invitrogen), PerCP-Cy5.5-anti-CD11c (Clone: N418, Invitrogen), eFluor-450-anti-Ki67 (Clone: SolA15, Invitrogen), Super Bright 645-anti-Sca-1 (Clone: D7, Invitrogen), FITC-anti-CD90.2 (Clone: 53-2.1, Invitrogen), APC-eFluor-780-anti-CD24 (Clone: M1/69, Invitrogen), APC-anti-rabbit IgG (H+L) (#31984, Invitrogen), anti-DCLK1 (ab31704, Abcam). Dead cells were excluded with the Fixable Aqua Dead Cell Stain Kit (Invitrogen). The BD Fix/Perm kit was used for intracellular staining. Samples were acquired on the BD LSRFortessa and analyzed with FlowJo Software (Treestar).

### Murine and human organoid cultures

Murine organoids were generated from crypts isolated from ileum as previously described.^25, 56^ Briefly, the ileum was opened, cleaned, scraped to remove villi, and cut into 1-cm pieces. Tissue was incubated in chelation buffer (2 mM EDTA in PBS) for 30-min at 4°C with rotation then transferred into shaking buffer (PBS, 43.3 mM sucrose, 54.9 mM sorbitol) and shaken by hand. Crypts were plated in Matrigel (Corning) with organoid culture media (60% Advanced DMEM/F12 supplemented with 10 mM HEPES, 2 mM L-glutamate, 40% L-WRN conditioned media, 1x N2, 1x B27, 50-ng/mL murine EGF, and 10 µM Y-27632) overlaid. Media was changed every 2-3 days.

For human organoids, de-identified terminal ileal biopsies were obtained from pediatric patients undergoing endoscopic analyses. Donors were recruited prospectively as controls for participation in the IBD Biorepository protocol at CCHMC (IRB 2011-2285) and histologic analyses confirmed patients as non-IBD healthy controls. Ileal biopsies were first incubated in strip buffer (PBS, 5% FBS, 1 mM EDTA, 1 mM DTT), then incubated in wash buffer (Advanced DMEM, 10% FBS, 2 mM L-glutamate, penicillin-streptomycin) with 2 mg/mL collagenase type I (Invitrogen) at 37°C for 5-15 mins with vigorous mixing as previously described.^57^ Crypts were plated in Matrigel with Human Organoid Growth Media (StemCell). Once organoids were established, they were maintained in human organoid culture media (50% Advanced DMEM/F12 supplemented with 10 mM HEPES, 2 mM L-glutamate, 50% L-WRN conditioned media, 1x N2, 1x B27, 50-ng/mL human EGF, and 10µM Y-27632). Organoids were stimulated with 1 mM butyrate (murine) or 0.5 mM butyrate (human) for 24 hrs, then treated with 20 ng/mL of IL-13 (human or mouse, PeproTech) for 72 hrs. For HDAC3 deletion in organoids derived from HDAC3^ΔIEC-IND^ mice, cells were treated with 1 µM of hydroxytamoxifen (4-OHT, Sigma) for 24 hrs prior to other treatments. For HDAC3i treatments, human organoids were stimulated with 10 µM of RGFP966 for 24 hrs before IL-13 treatment. For immunostaining, human and mouse organoids were plated in 8-well u-slide (ibidi) and fixed in 4% paraformaldehyde for 20-min. Organoids were permeabilized for 20-min in PBS containing 0.5% Triton X-100 followed by a 30-min incubation in blocking buffer (PBS, 0.2% Triton X-100, 0.05% Tween, 1% BSA, and 1% normal goat serum). Organoids were stained overnight at 4°C in blocking buffer containing DAPI (Sigma), AlexaFluor-647-Phalloidin (Invitrogen) and anti-DCLK1 (ab31704, Abcam) for mouse or AlexaFluor-488-anti-pEGFR (ab205827, Abcam) for human. Organoids were washed and imaged using a confocal microscope (Nikon).

### Gene expression and RNA-sequencing

RNA from primary IECs and organoids were isolated using the RNeasy Kit (Qiagen) following manufacturer’s protocol. cDNA was synthesized using the Verso reverse transcriptase kit (Thermo Fisher) following the manufacturer’s protocol. Real-time PCR was performed using SYBR green (Applied Biosystems) and analyzed using the following murine primers: mHPRT forward 5’- GATTAGCGATGAACCAGGT-3’, mHPRT reverse 5’- CCTCCCATCTCCTTCATGACA-3’, mPou2F3 forward 5’- TGGTGAATCTGGAGCCCATGC-3’, mPou2F3 reverse 5’- AGAGTCACCCAGATCCTCCG-3’, mDCLK1 forward 5’- TGAACAAGAAGACGGCTCACTCC-3’, mDCLK1 reverse 5’- GCTGGTGGGTGATGGACTTGG-3’, mIL-25 forward 5’- ACAGGGACTTGAATCGGGTC-3’, mIL-25 reverse 5’-TGGTAAAGTGGGACGGAGTTG-3’, mSprouty2 forward 5’- TCCACCGATTGCTTGGAAGT -3’, mSprouty2 reverse 5’- CCATCAGGTCTTGGCAGTGT -3’. Human cells were analyzed with the following human primers: hHPRT forward 5’- GATTAGTGATGATGAACCAGGTT-3’, hHPRT reverse 5’- CCTCCCATCTCCTTCATCACA-3’, hTRPM5 forward 5’- GTGTCTGGAGTCACAGATCAACTA-3’, hTRPM5 reverse 5’- GCTCAGGTGTCCGAGGGA-3’, hPou2F3 forward 5’- GGCAGGATGGTGAATCTGGAG-3’, hPou2F3 reverse 5’- <colcnt=4> GACATGGGCTGGCATTTAGC-3’, hSprouty2 forward 5’- CCCCTCTGTCCAGATCCATA-3’, hSprouty2 reverse 5’- CCCAAATCTTCCTTGCTCAG-3’. For global expression analyses, RNA was isolated from organoids and subjected to Illumina next-generation sequencing. Sequencing reads were trimmed to remove bar codes and mapped to the mouse genome (GRCm38) using Bowtie2.^58^ Differential expression analysis was performed using DESeq2 within Seqmonk (V1.47.1) (p<0.05, fold change >1.5).^59^

### Chromatin-immunoprecipitation

ChIP-qPCR on IECs was performed as described previously.^24, 25, 60^ Briefly, cells were fixed for 10 min in 1% PFA at room temperature, followed by quenching with 125 mM glycine for 10 min. After a two-step wash with cold PBS, fixed cells were lysed, and nuclear extracts were washed in TE 0.1% SDS with protease inhibitors and sonicated using a S220 Focused-ultrasonicator (Covaris). Prior to immunoprecipitation, sheared chromatin was precleared for 20 min at 4°C using Protein G Dynabeads (Thermo Fisher Scientific). Immunoprecipitations were performed using fresh beads and anti-Histone H3 acetyl K9 (H3K9Ac) antibody (#06-942, Millipore) using a SX-8G IP-STAR automated system (Diagenode) with the following wash buffers: (1) RIPA 150 mM NaCl, (2) RIPA 400 mM NaCl, (3) Sarkosyl Buffer (2 mM EDTA, 50 mM Tris-HCl, 0.2% sarkosyl (Lauroylsarcosine sodium salt (Sigma)), and (4) TE 0.2% Triton X-100. Immunoprecipitated chromatin were treated with Proteinase K (Thermo Fisher Scientific) at 42°C for 30 min, 65°C for 4 hr, and 15°C for 10 min in elution buffer (TE 250 mM NaCl 0.3% SDS). Phenol:chloroform isoamyl alcohol with Tris-HCl (pH 8.0) and chloroform phase-separation were used to isolate DNA, followed by overnight ethanol precipitation for primary IECs or via QIAquick PCR purification kit (Qiagen) for organoids. ChIP DNA was treated with molecular grade RNase and subjected to quantitative real-time PCR using the following custom-made primer pairs; murine Sprouty2 promoter forward 5’- ATGAACAAGAGGCCGAAGCG-3’ and murine Sprouty2 promoter reverse 5’- CTCCACTTGGGAGCAGCC-3’; or human Sprouty2 promoter forward 5’- AAGACTAATCTGGGCCACGC-3’ and human Sprouty2 promoter reverse 5’- TCCGCAAAGGGGAACAGAAA-3’. Data were analyzed as percent enrichment over input DNA.

### Tissue immunofluorescence

Ileal biopsies were fixed in 4% paraformaldehyde (PFA), embedded in paraffin, and sectioned. Sections were deparaffinized, rehydrated, and heated in antigen retrieval buffer (Tris-EDTA buffer with 0.05% Tween-20 at pH 9.0 or sodium citrate buffer at pH 6.0) for 20 minutes using a water bath at 95°C. Sections were washed in PBS and blocked in PBS containing 0.1% Triton X-100 and 5% normal donkey serum (NDS) (Jackson ImmunoResearch) for 45 minutes at room temperature. Sections were incubated with primary antibodies overnight at 4°C, washed three times in PBS containing 0.05% Tween-20, and then incubated with secondary antibodies for 1 hour at room temperature. To visualize the nuclei, sections were incubated in PBS containing DAPI for 10 minutes at room temperature. Stained sections were washed three times in PBS containing 0.05% Tween-20 before mounting with ProLong Gold Antifade Mountant (ThermoFisher). Immunofluorescence was performed using the following primary antibodies: mouse anti-E-Cadherin (Clone: 36/E-Cadherin, BD Biosciences), rabbit anti- DCLK1 (ab31704, Abcam), and goat anti-TdTomato (AB8181-200, Origene). The following secondary antibodies were all from Jackson ImmunoResearch unless noted: AlexaFluor-647-donkey anti-mouse IgG (H+L) (715-605-151), AlexaFluor-488-donkey anti-rabbit IgG (H+L) (711-545-152), AlexaFluor-594-donkey anti-goat IgG (H+L) (705-585-147), and AlexaFluor-555-donkey anti-mouse IgG (H+L) (ab150106, Abcam). Images were acquired with a Nikon A1 inverted confocal microscope. Images were processed using Nikon NIS-elements software and Fiji: ImageJ.

### Quantification of short-chain fatty acids

Fecal samples were collected and flash frozen. Samples were prepared for gas-chromatology mass spectrometry as previously described.^61^ Briefly, fecal samples were homogenized with water at a concentration of 1 mg of sample per 10 µL of water. Samples were sonicated for 15 mins, then centrifuged at top-speed at 4°C for 15 mins. Buffer and derivatization reagents were added in supernatant following the ratio of 5:2:14 (Supernatant: Buffer: derivatization reagent, v/v/v). 100 µL of supernatant was transferred to a glass vial and 40 µL PBS (buffer) and 280 uL of pentafluorobenzyl bromide (derivatization reagent) was added. Samples were vortexed for 2 mins and incubated in a water bath at 60°C for 90 mins. 100 µL of hexane was added after the sample cooled down to extract short chain fatty acids. Samples were vortexed, centrifuged, then the top layer (hexane phase) was transferred to a gas chromatography vial for GC-MS analysis. Sample preparation and GC-MS analysis was conducted by Center for Regulatory and Environmental Analytical Metabolomics (CREAM) at University of Louisville.

### Statistical analysis

Results are expressed as mean ± SEM. Statistical significance was determined with the Student’s t-test or one-way ANOVA. Results were considered significant at *p < 0.05; **p < 0.01; ***p < 0.001, ****p < 0.0001. Statistical significance was calculated using Prism version 7.0 (GraphPad Software).

## Supplemental Data Figure Legends

**Figure S1. Stem cell-intrinsic HDAC3 drives tuft cell hyperplasia. (A)** Fluorescence staining of tuft cells (DCLK1^+^, green) in ileum, **(B)** frequency of DCLK1^+^ tuft cells, and **(C)** frequency of ILC2s by flow cytometry in HDAC3^ΔSTEM^ mice treated with succinate then vehicle or tamoxifen. ILC2s gated Live, CD45^+^, lineage (CD4, CD8a, CD11b, CD11c, B220, Ly6G)^-^, CD90.2^+^, CD127^+^, Sca-1^+^, KLRG1^+^. Scale bars, 50µM. Data are representative from at least three independent experiments, 3-4 mice per group. *p<0.05, **p<0.01, ***p<0.001, ****p<0.0001, ns=not significant.

**Figure S2. Microbiota control tuft cell differentiation and type 2 immunity through epigenetic regulation of intestinal stem cells.** The epigenetic modifying enzyme, HDAC3, promotes tuft cell expansion and downstream tuft cell-induced type 2 immune responses during basal conditions and helminth infection. Microbiota-derived butyrate blocks tuft cell differentiation by inhibiting HDAC3 activity in intestinal stem cells. These data reveal an active role for microbiota-derived metabolites in epigenetically limiting tuft cell differentiation to control type 2 immunity. This figure was created using Biorender.com.

