## Supplementary Figures for "Microbiota epigenetically direct tuft cell differentiation to control type 2 immunity"

**Figure S1. Stem cell-intrinsic HDAC3 drives tuft cell hyperplasia.**

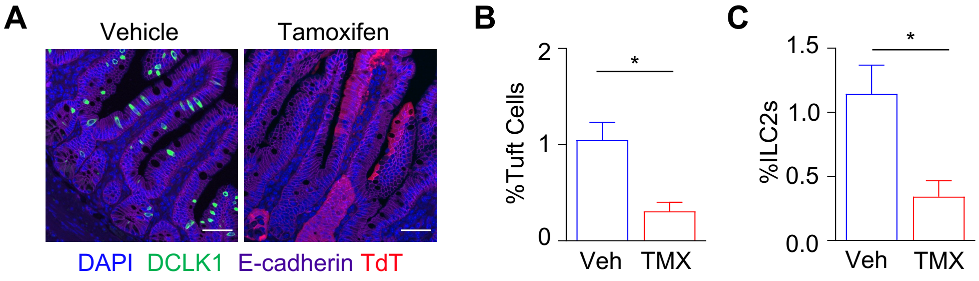


**Figure S1. Stem cell-intrinsic HDAC3 drives tuft cell hyperplasia. (A)** Fluorescence staining of tuft cells (DCLK1$^+$, green) in ileum, **(B)** frequency of DCLK1$^+$ tuft cells, and **(C)** frequency of ILC2s by flow cytometry in HDAC3$^{\Delta STEM}$ mice treated with succinate then vehicle or tamoxifen. ILC2s gated Live, CD45$^+$, lineage (CD4, CD8a, CD11b, CD11c, B220, Ly6G)$^-$, CD90.2$^+$, CD127$^+$, Sca-1$^+$, KLRG1$^+$. Scale bars, 50μM. Data are representative from at least three independent experiments, 3-4 mice per group. *p<0.05, **p<0.01, ***p<0.001, ****p<0.0001, ns=not significant.

**Figure S2. Microbiota control tuft cell differentiation and type 2 immunity through epigenetic regulation of intestinal stem cells.**

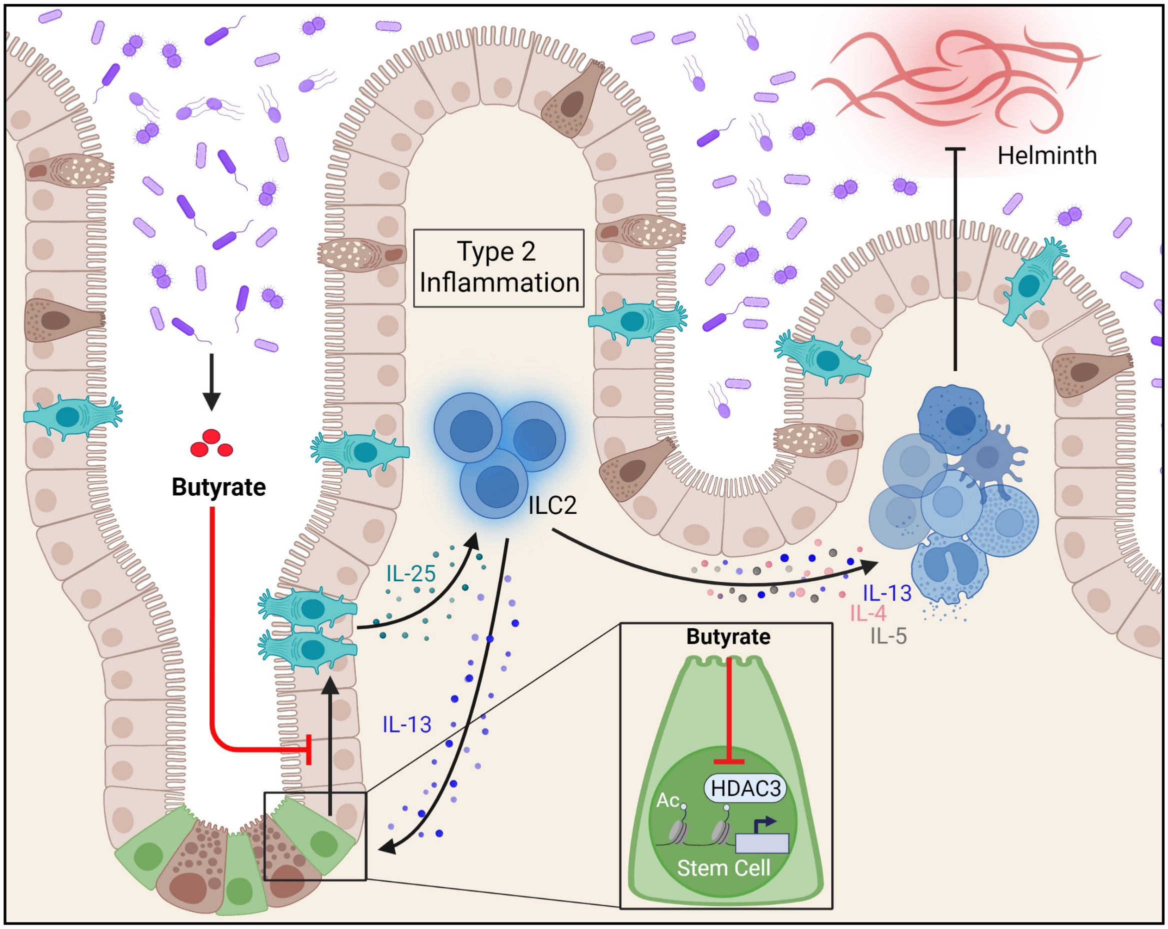


**Figure S2. Microbiota control tuft cell differentiation and type 2 immunity through epigenetic regulation of intestinal stem cells.** The epigenetic modifying enzyme, HDAC3, promotes tuft cell expansion and downstream tuft cell-induced type 2 immune responses during basal conditions and helminth infection. Microbiota-derived butyrate blocks tuft cell differentiation by inhibiting HDAC3 activity in intestinal stem cells. These data reveal an active role for microbiota-derived metabolites in epigenetically limiting tuft cell differentiation to control type 2 immunity. This figure was created using Biorender.com.
